# A Mathematical Model of Exploration and Exploitation in Natural Scene Viewing

**DOI:** 10.1101/2020.04.16.044677

**Authors:** Noa Malem-Shinitski, Manfred Opper, Sebastian Reich, Lisa Schwetlick, Stefan A. Seelig, Ralf Engbert

**Affiliations:** Institute of Mathematics, University of Potsdam, Potsdam, Germany; Department of Artificial Intelligence, Technische Universität Berlin, Berlin, Germany; Department of Psychology, University of Potsdam, Potsdam, Germany

## Abstract

Understanding the decision process underlying gaze control is an important question in cognitive neuroscience with applications in diverse fields ranging from psychology to computer vision. The decision for choosing an upcoming saccade target can be framed as a dilemma: Should the observer further exploit the information near the current gaze position or continue with exploration of other patches within the given scene? While several models attempt to describe the dynamics of saccade target selection, none of them explicitly addresses the underlying Exploration–Exploitation dilemma. Here we propose and investigate a mathematical model motivated by the Exploration–Exploitation dilemma in scene viewing. The model is derived from a minimal set of assumptions that generates realistic eye movement behavior. We implemented a Bayesian approach for model parameter inference based on the model’s likelihood function. In order to simplify the inference, we applied data augmentation methods that allowed the use of conjugate priors and the construction of an efficient Gibbs sampler. This approach turned out to be numerically efficient and permitted fitting interindividual differences in saccade statistics. Thus, the main contribution of our modeling approach is two–fold; first, we propose a new model for saccade generation in scene viewing. Second, we demonstrate the use of novel methods from Bayesian inference in the field of scan path modeling.

**Author summary:** The Exploration–Exploitation dilemma is general concept that has been investigated in human information processing. We investigate whether the Exploration–Exploitation trade–off is a viable approach to model sequences of fixations generated by a human observer in a free viewing task with natural scenes. Variants of the basic model are used to predict to the experimental data based on Bayesian inference. Results indicate a high predictive power for both aggregated data and individual differences across observers. The combination of a novel model with state-of-the-art Bayesian methods lends support to the Exploration–Exploitation framework in the field of eye-movement research.

## Introduction

The human visual system acquires high-acuity information from a rather small region (the fovea) surrounding the center of gaze [1]. The foveal organization of the visual system has two immediate consequences. First, visual perception of natural scenes depends critically on the control of precise and fast eye movements (saccades) that move regions of interest into the fovea for high-acuity processing. During a typical visual task (e.g., scene viewing or reading), saccades occur at a rate of 3 to 4 per second [2]. Second, the decision process for an upcoming saccade target poses a dilemma: should the observer further exploit the information near the fovea or continue with exploration of other patches within the given scene? The latter problem is critical for scene viewing [3, 4] and relevant to the broader field of cognitive processes in knowledge acquisition [5].

Observers select saccade targets from a priority map [6] that represents objects and regions within a given scene according to their attentional weight. Over the last decades, computational modeling of visual attention for natural scenes [7] resulted in a broad range of successful models [8] of priority maps. These models use feature maps to combine low-level saliency and top-down control. Recently, deep neural network (DNN) models achieved state–of–the–art performances in predicting saliency maps from images [9,10]. From these advances, the problem of modeling priority maps seems basically solved [11]: for an arbitrary natural image, computational models can generate a prediction of fixation density in experiments with human observers.

The next step in modeling human visual behavior is fundamentally related to the fact that eye movements introduce sequential steps in information processing. Since access to visual information is effectively limited to the fovea, the full sequence of saccadic gaze shifts (scan path) needs to be modeled in order to understand the underlying principles. Understanding how human observers shift their attention while looking at an image requires quantifying the scan paths (Figure1).

**Fig 1.**
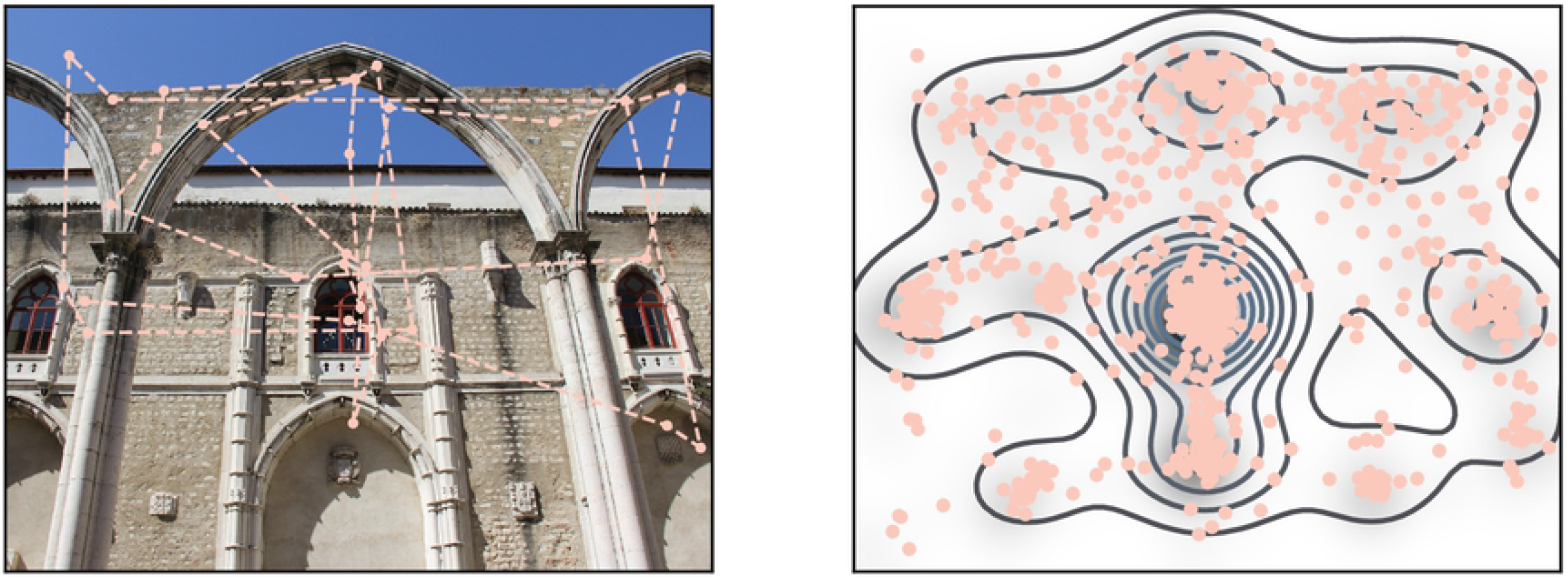
Experimental scanpath and fixations density. *Left*. An image and a scan path. Each dot is a fixation and the dashed line illustrates the saccade. *Right*. The empirical fixation density map as generated by aggregating the fixations from all subjects for a given image.

So far, few models for scan path generation and prediction have been proposed. These models can be generally classified into two groups, where one group of models is hypothesis–based and the other is hypothesis–free. The second group includes models which use state of the art deep learning techniques [12], [13]. While these models capture structure present in the data, they provide only very limited insights into the underlying principles of scan path generation. Another critical point for experimental research is that deep learning models require a lot of training data, which are typically unavailable for single observers. Thus, current deep learning approaches do not capture interindividual differences in statistical properties of scan paths.

Hypothesis–based models rely on cognitive and neural assumptions of human perception and oculomotor control that were derived from known biological mechanism and well-established experimental effects [14–17]. Thus, the key goals of parametric models are (i) to implement these assumptions in a fully quantitative way and build a generative model, (ii) to fit the model to experimental data for hypothesis testing (statistical inference), and, finally, (iii) to provide explanations for interindividual differences in experimental data sets [18].

In the current study, we introduce a new model which belongs to the class of parametric models. Our central hypothesis is that the generation of scan paths can be phrased in the context of the Exploration–Exploitation trade-off. In this view, the generation of each saccade is a decision process, where the observer has to choose between a short saccade for staying in the immediate surrounding of the current fixation (exploitation) and a long saccade to explore a new region of the visual environment (exploration). We assume that the decision is based on the information currently available to the observer. Specifically, this assumption translates into a higher probability of an exploitation saccade if the ratio of priority values of the current fixated location and the previously fixated location is high. This hypothesis follows an assumption that the area next to a location with high priority also has high priority. This assumption is valid for natural images used in this work.

The idea of exploration and exploitation intentions in visual behavior was studied previously by Gameiro et al. [3]. Their work demonstrated experimentally that the tendency for exploration or exploitation, measured by saccade amplitude and fixation duration, depends on size and spatial properties of the stimulus. The characterization of the exploratory or exploitative tendencies was done using the statistics of the entire scan paths.

Different from the approach taken by Gameiro et al. [3], we use the Exploration–Exploitation terminology to analyze individual saccades rather than entire scan paths. Our generative model tags each saccade as either an exploration or an exploitation step. We aim at a minimal model to keep computations efficient and to facilitate interpretations of the model behavior. A critical component of our approach is the application of Bayesian statistics to fit the model to experimental scan path data. We use the fitted model to quantify how well the model describes the experimental data. Further we test different variations of the model, which correspond to different hypotheses, to determine which hypothesis corresponds best to the experimental data.

In the next section we describe the details of our basic model and explain the computation of the likelihood function as a fundamental tool for statistical inference. We construct the model in a modular way and relate each part to one of the assumptions we would like to investigate. Next, we describe the process of fitting the model parameters to experimental data. In the Results, we compare several statistics of simulated data to the statistics of the experimental data. We also analyze different variants of the basic model and quantify how well each one of them describes the data using the model’s likelihood function. We close with the Discussion of our results in the context of current problems in understanding scan path generation during scene viewing.

## Materials and methods

### The Exploration–Exploitation Model for scan path generation

Our theoretical investigation of exploration and exploitation in saccadic behavior is based on the implementation of a probabilistic generative model. The static viewer independent priority map for saccadic selection [6] is thought to be the combined result of early visual processing or *saliency* [7] and top-down cognitive control. While various models for the computation of static priority maps exist, we extend the modeling approach to the generation of scan paths for a given static saliency map. For simplicity, we use the time-averaged fixation density [16] as an approximation of the saliency of a given image.

The static saliency map is a function *s*(*z*): ℝ^2^ ↦ ℝ^+^ with *z* = (*x,y*) being a location in an image and *s*(*z*) being the probability of an average viewer to fixate this location (its saliency). As mentioned above, we approximate the saliency map by the experimentally-observed fixation density and we use *s*(*z*) or *s_z_* to refer to the saliency map or the fixation density of the image at location *z*.

Generally, scan paths are sequences of fixation locations and fixations duration. In this work we model only the spatial properties of gaze control. We account only for the temporal ordering of the fixations and do not model the fixation duration. In these settings, a scan path is written down as *Z* = {*z_1_, z_2_,…, z_t_,…, z_T_*} with *T* being the number of fixations in the scan path and *z_t_* being the location of the tth fixation.

We begin constructing our model by assuming that the saccade generation process is a second order Markov process, which means that the probability *p*(*z* = *z_t_*) of fixating on a specific location *z_t_* at time step *t* depends only on the location of the fixation at time *t* − 1 and the fixation at time *t* − 2. The probability of a full scan path is written as

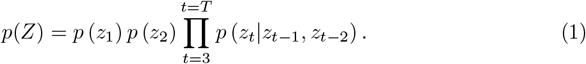

The choice of the second order Markov process reflects our hypothesis regarding the scan path generation and will become clear in the upcoming paragraphs. In principle, it is possible to construct a simpler model which corresponds to first order Markov process. This would correspond to slightly different assumptions regarding the scan path generation and we refer to such a model in the section discussing simplified models.

We describe the probability of the next fixation being *z_t_* given that the previous two fixation location were *z_t-1_* and *z_t-2_* in terms of competing exploitation and exploration policies:

#### Exploitation

The next fixation location is chosen close to the current fixation location following a Gaussian distribution around the current fixation location with covariance e, normalized over the entire image. This can be written as

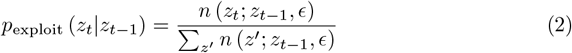

where *n*(*z_t_; z_t-1_, ϵ*) is a Gaussian density with mean *z_t-1_* and covariance 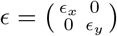.

#### Exploration

A potential exploration policy is that the next fixation location is chosen randomly from the static saliency map of the image. This policy leads to very large saccade amplitudes which are known to be less probable [19]. To integrate this prior regarding the saccade amplitudes knowledge into the model - instead of choosing the next fixation location from the the saliency map, we modulate the saliency map by a Gaussian distribution, which gives a higher weight to areas of high saliency which are closer to the current location.

This approach results in the following expression for the exploration policy

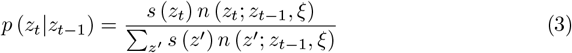

with *ξ* a diagonal covariance matrix similarly to *ϵ*, *ξ_x_* > *ϵ_x_* and *ξ_y_* > *ϵ_y_*.

Having Equation (3) as an exploration policy may result in short saccades similar to the ones generated by the exploitation policy when the current fixation is in a high priority area. A solution is to create a repulsion mechanism that forces the saccades generating by the exploration policy to be of at least a certain length. This is achieved by the following expression

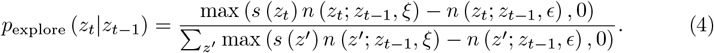

To avoid negative values for the likelihood we take the maximum between the subtraction and 0. Figure2 visualizes the two distributions formulated in Equations (2) and (3).

**Fig 2.**
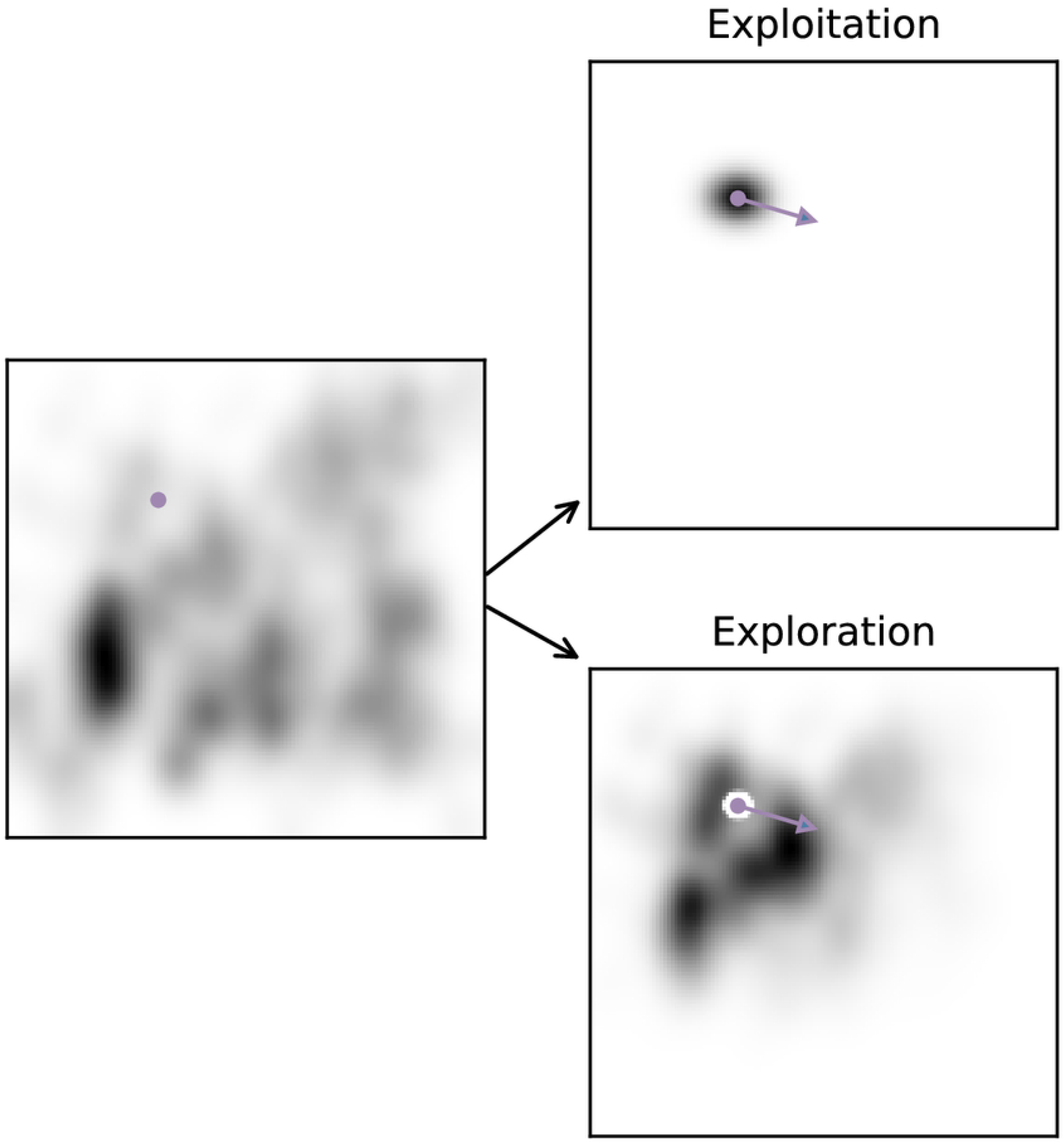
Exploration and exploitation policies. On the left is an example of an empirical saliency map, the dot indicates a fixation location. On the right are the probability maps generated by either the exploitation policy (upper panel) or the exploration policy (lower panel). The arrow indicates a saccade.

Our assumption is that each fixation comes either from the exploration policy described in Equation (4) or the exploitation policy described in Equation (2). This can be represented as a mixture model

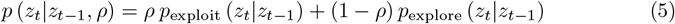

The model parameter *ρ* describes the tendency to perform either an exploration or exploitation step. It can be fixed based on prior knowledge or inferred from the experimental data. The condition *ρ* > 0.5 implies that the probability for an exploitation step is larger than for an exploration step for every saccade.

Next we include in our model the assumption that *ρ* changes depending on the fixation location. We use the notation *ρ_t_* to indicate that the fixation *z_t_* was generated based on *ρ_t_*. Importantly, this notation does not imply that *ρ_t_* is necessarily a function of *z_t_*.

We assume that the decision whether to make an exploration or an exploitation step depends on the ratio between the priority values of the current and previous fixated locations. The result is that the viewer is more likely to make an exploitation step if the saliency value of the current fixated location is higher than the saliency value of the previous fixated location. We include this in the model with the following expression for *ρ_t_*

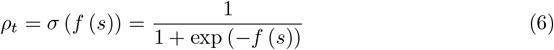

with

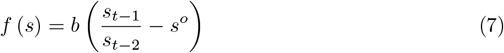

with *s_t-1_* = *s*(*z_t-1_*) and *b* and *s°* being scalar variables.

Combining Equation (1) and Equation (5), the model likelihood is written as

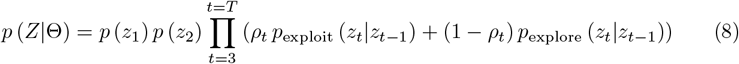

with model variables Θ = {*ϵ, ξ, b, s°*}. Here, we chose to sample the first and second fixation from the empirical static saliency map such that *p*(*z*) = *s*(*z*).

Figure3 presents a scan path generated by our model given a particular saliency map, along side a scan path recorded experimentally from a viewer viewing the image corresponding to the saliency map.

**Fig 3.**
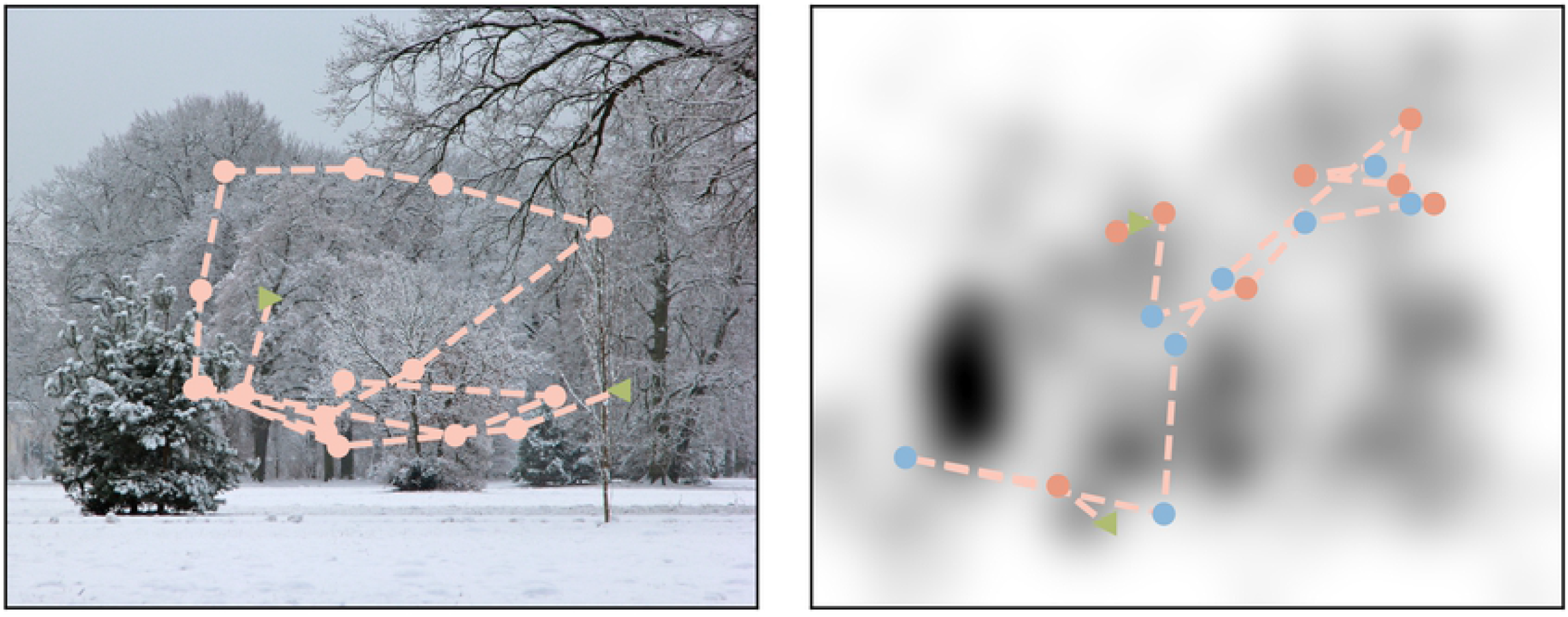
Experimental and simulated scan paths. *Left*. An image and a scan path recorded from a human observer. *Right*. The experimental static saliency map and a scan path generated by the Exploration–Exploitation Model. The green arrow pointing right represents the first randomly selected fixation location. The green arrow pointing left represents the last fixation in the scan path. The blue dots are fixations that were generated from an exploration step and the pink dots are fixations that were generated from an exploitation step.

### Simplified models

To test the different assumptions behind our full model described above, we construct three simpler models and compare their performances to the performance of the full model in the Results. To construct the models we remove one by one the assumptions on which the model is based. This results in the following competing models:

#### Local Choice Model

Equation (6) describes the assumption that the decision between an exploration and an exploitation step depends on the ratio between the priority value of the current fixation location and the priority value of the previous fixation location. A competing assumption would be that the decision depends only on the priority value of the current fixation location. In this case we keep the model the same and only change *f*(*s*)

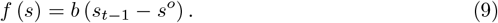

#### Fixed Choice Model

We test the assumption that the decision between the policies does not depend on the saliency value of previous fixation. In this simplification of the model, rather than having *ρ_t_* = *f*(*z_t-1_, z_t-2_*) we have a fixed probability to chose each policy with *ρ_t_* = *ρ*.

#### Local Saliency Model

Last, we challenge the approach of two competing policies. In this variation of the model, each fixation is generated from a modulation of the empirical saliency map with a Gaussian around the current fixation location. This corresponds to the following fixation location likelihood

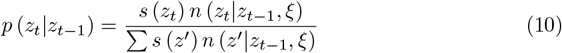

In the next section we describe the inference process of the full Exploration–Exploitation Model. As the three models described above are simplified versions of the full model we do not describe their corresponding inference processes as they can be easily derived from the inference of the full model.

### The inference process

Our approach is based on experimental results and we derive the model parameters from observed data in a Bayesian framework. This approach allows us to include prior knowledge regarding the different model parameters based on known spatial features of scan paths. It also allows us to obtain distributions over the model parameters, rather than point estimates, and to compare different variations of the model via the respective test–data likelihoods.

In the previous section we defined the likelihood of the data. Next, we describe the data augmentation methods which allow us to identify conjugate priors and construct an efficient Gibbs sampler [20] using the full conditional distributions over the model parameters.

The idea behind data augmentation [21] is adding latent variables to the model, which can be considered as unobserved data, in a way that simplifies the inference of the parameters of interest. We use the standard approach and augment the Exploration–Exploitation likelihood by

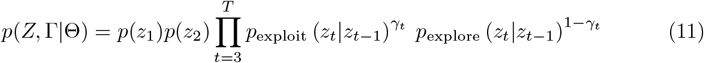

with

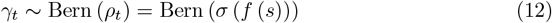

and marginalizing over ⌈ results in Equation (8).

The augmentation defines a modified generative process for the model. At each time step a variable *γ_t_* is drawn from a Bernoulli distribution with bias *ρ_t_*. If the result is 1then the next saccade is generated following the Exploitation policy. If the result is 0, the saccade is generated from the exploration policy. This construction reflects our assumption regarding the cognitive process underlying scan path generation, where each saccade follows either the exploration or the exploitation policy.

For a simple two–component mixture model with normal distribution, the augmentation described above would have been sufficient for the derivation of a Gibbs sampler [22]. As the model we constructed is more complex, we need to handle the sigmoid link–function in Equation (6) and the non-trivial form of the Exploration distribution in Equation (3).

With the Sigmoid function in Equation (6) there is no straightforward way to define conjugate priors for the parameters *b* and *s°* which are needed for a Gibbs sampler. To achieve conditional probabilities which are easy to sample from, we augment the model with another set of latent variables *w_t_*, which follow a Pólya-Gamma distribution

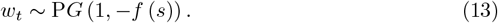

As described in [23] for the case of logistic regression, the usage of this augmentation scheme results in conditional distributions for *b* and *s°* which are Gaussian and can be sampled from easily. The full derivation of the discussed conditional distributions can be found in the supplementary material.

After adding the two sets of latent variable we can define conjugate priors for the parameters:

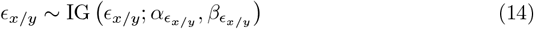

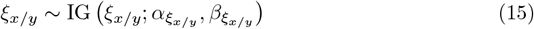

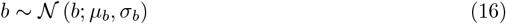

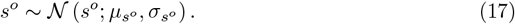

The prior distributions described above include hyperparameters. These parameters were chosen and not inferred from the data. The hyperparameters related to the prior distributions over *ϵ_x/y_* and *ξ_x/y_* were chosen based on known characteristics of human saccades. The hyperparameters related to *b* and *s°* were chosen to be on the same scale of the average 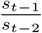 from the data. Further, all of the hyperparameters were chosen to induce wide prior distributions.

Combining the likelihood in Equation (8) with the priors defined above, the posterior distribution over the model parameters and the latent parameters is given by

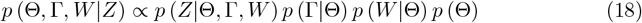

with

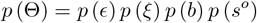

We can sample easily from the conditional distributions of *b* and *s°*. This is not the case *ϵ* for *ξ* and because of the form of p_explore_.

Due to the complex form of the exploration expression in Equation (4), which includes both *ξ* and *ϵ*, there is no closed form for the conditional distribution of these parameters. Thus, we resort to a technique known as MCMC within Gibbs [24,25] and in each iteration of the Gibbs sampler we evaluate the conditional distributions of *ξ* and *ϵ* using an Hybrid Monte Carlo step [26], also known as Hamiltonian Monte Carlo.

For further technical details regarding the augmentation and the HMC sampler please see the supplementary material.

## Results

In this work we propose an Exploration–Exploitation Model for scan path generation. In the previous section we derived the model equations from the basic Exploration–Exploitation approach. We described the inference process of our model when applied to experimental data. In this Section, we present the results of the inference process. First, we analyzed the reliability of our procedures by fitting the model to artificial data generated from the model with known parameter values. Next, we fit the model to the experimental data and test the statistics of the data generated from the model against the experimental data. Finally, we compare different versions of the model.

### Model parameters estimation

As presented in the Methods Section, the inference process includes using an MCMC approach to evaluate the posterior function over the model parameters. This approach is exact in the limit of an infinite number of samples but as we can only use a finite number of samples our the result is an approximation of the actual posterior. The distribution of the inferred parameters should concentrate around their real values.

When fitting the model to experimental data it is impossible to know the real values of the model parameters as they do not relate directly to any measurable features of the data. Thus, in order to assess the performance of the inference we use data simulated by the model, in which case we know the exact values used to generate the data. If the inference process is correct we expect the resulting posterior distribution to be concentrated around the ground truth values.

We generated data from our model with the parameter values that were inferred from the experimental data. In order to see whether the inference process will have reasonable results when fitting the experimental data, the size of the generated data set is comparable to the size of the experimental data for one subject.

Figure4 presents the distribution over model parameters as results from the inference process with data generated by the model. Each of the ten colored curves represents a different inference process started at a different point. As expected all the curves from different runs are similar in shape. The black dashed curves present the prior distribution over the parameters. To test the model we chose the prior distribution so their modes do not overlap with the values used in the data generation. As expected the mode of the inferred parameter distribution is close to the real values used in the data generation which are noted by the vertical solid line.

We tested the model on generated data that have similar properties to the experimental data. Generated scan paths had lengths similar to the lengths of scan paths recorded experimentally. This could have the result that the generated data does not have sufficient information regarding the underlying model parameters and it explains the deviation of the distribution mode from the true parameter values.

**Fig 4.**
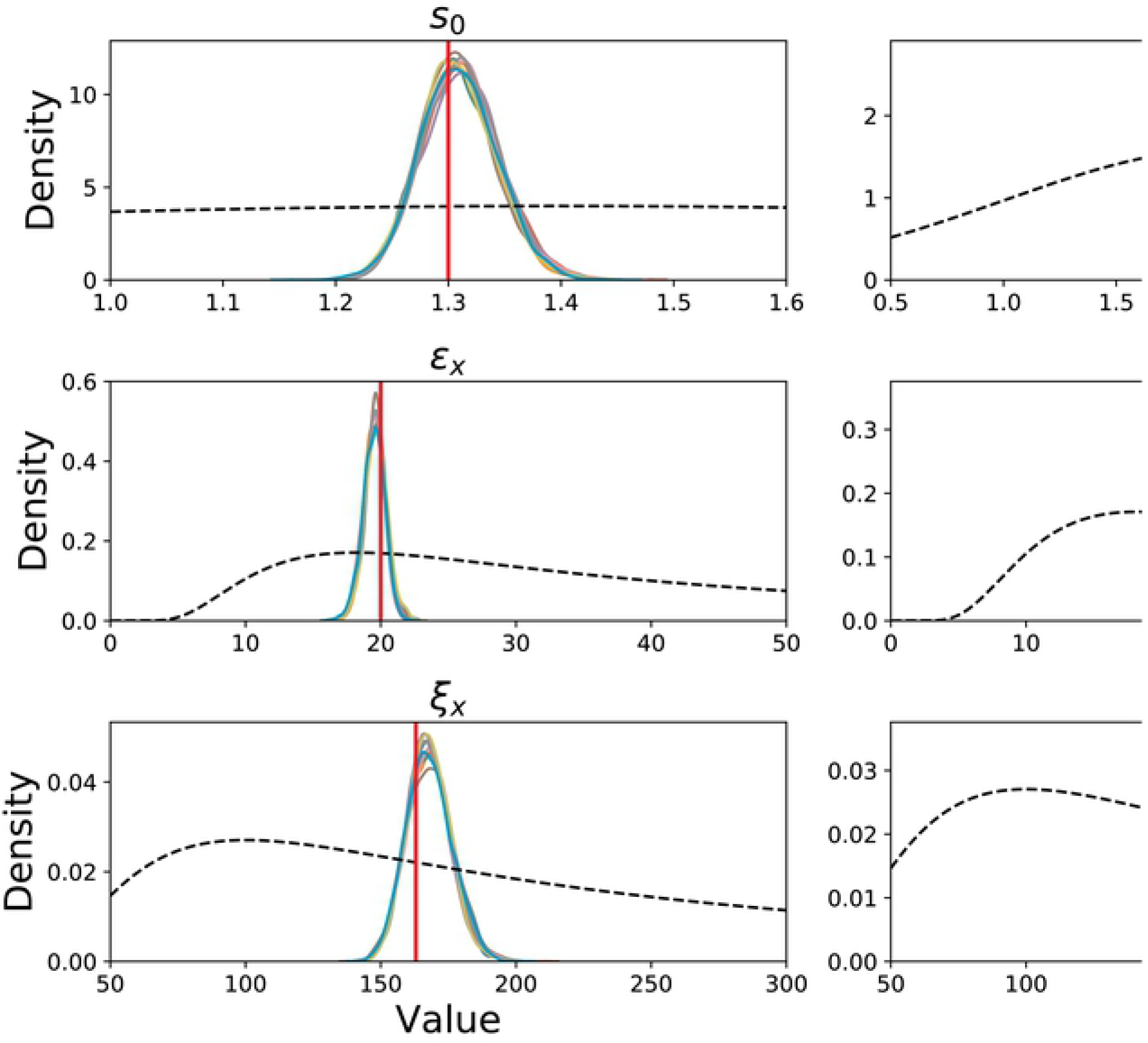
Inference results on simulated data. Model parameter recovery. To test the inference algorithm we fit the model to simulated data with known parameters values. Each panel includes the inferred posterior distribution of each parameter after the inference process. The ten curves present 10 different inference processes starting from different values. The vertical lines are the values with which the data was generated. The black dashed curve is the prior distribution. The plotted densities are not normalized.

### Model performance on experimental data

Our model was derived from a set of hypotheses regarding the cognitive process of saccade generation. In order to test the validity of the model, and of the corresponding hypotheses, we fit the model to the data, simulate new data using the model and check whether the features of the simulated data correspond to the features of the experimental data.

The data set used here includes the scan paths of thirty five human observers performing a memorization task over thirty natural images. The same data set was used before to evaluate other scan path models [16], [18]. The acquisition of the data was carried out in accordance with the Declaration of Helsinki, and informed consent was obtained for experimentation by all participants. Data from three subjects were excluded as the inference process did not converge.

We fit a separate model for each subject, while using the same prior hyperparameters for all fitted models. We want to test whether the model captures subjects’ tendencies that generalize over images. Thus, we split the model and use 70% of the trials as training data, with which the model parameters are fitted, and use the remaining 30% of the data as the test set. All the reported results in this section are obtained from the test data.

### Saccade amplitude

The Exploration–Exploitation Model was designed to capture the different saccade amplitudes generated by subjects while observing an image in a free viewing task. To estimate the model’s performance we compare the amplitudes of the empirical saccades with the amplitudes of the saccades generated by the model. The comparison is done both at a population level and for each subject separately.

Figure5 compares the empirical saccade amplitude density with the saccade amplitude density of the scan paths that were generated by the full Exploration–Exploitation Model and the simplified versions presented previously. The density presented is over the entire population of subject. The black curve presents the empirical data. The orange curve corresponds to data generated by the full Exploration–Exploitation Model, and the other curves correspond to the different simplified models.

**Fig 5.**
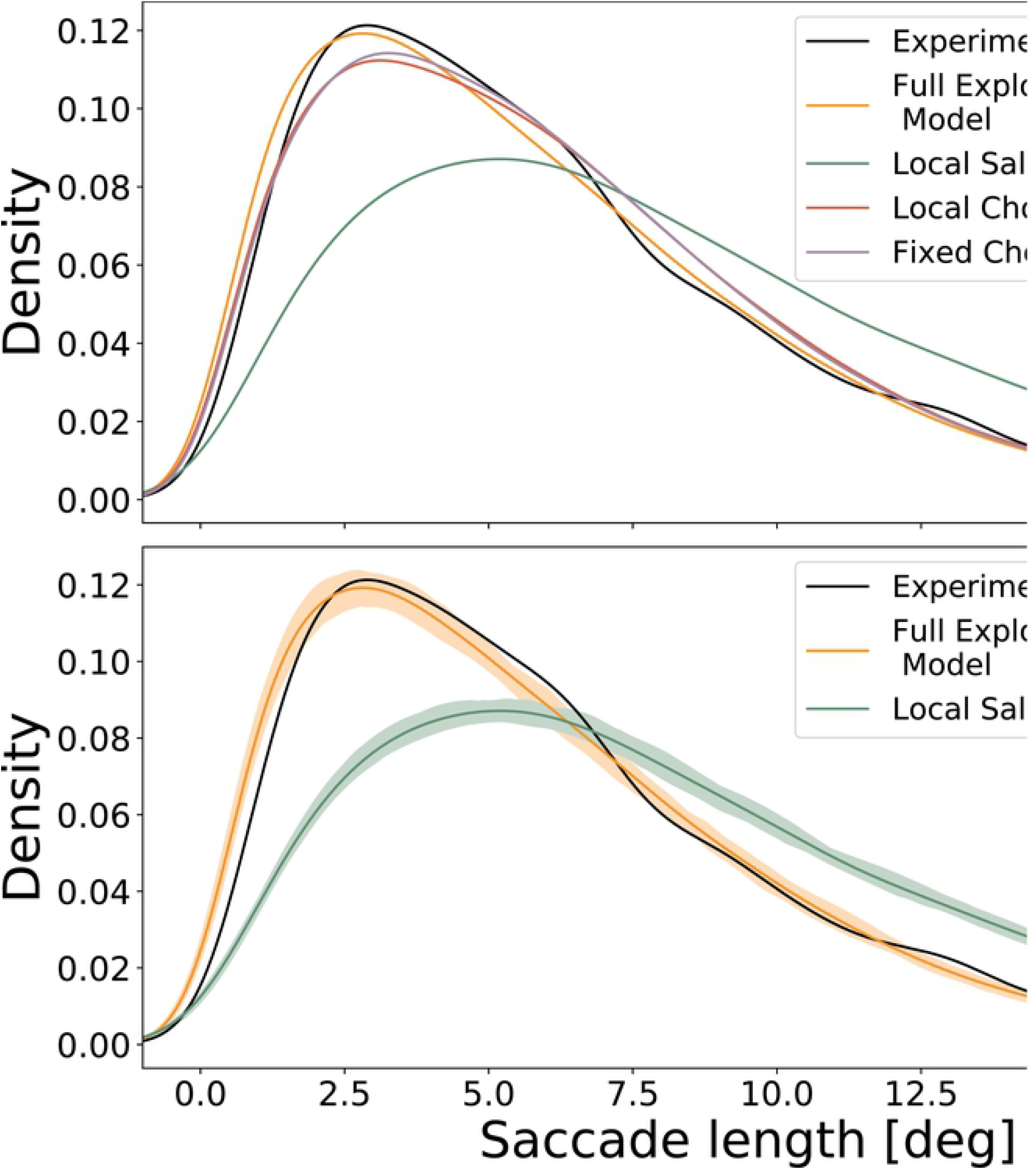
Saccade amplitude density - experimental and simulated. Saccade amplitude density, aggregated over the data from all participants, of the experimental data and data generated by the full model and the simplified competitor models. *Top*. Comparison of all models. *Bottom*. Comparison between the full Exploration–Exploitation Model and the Local Saliency Model. The shading corresponds to confidence bounds regarding the estimate of the model parameters. The full Exploration–Exploitation Model manages to capture the different kinds of saccade lengths, whereas the simpler models fail to do so.

Figure5 shows that the full Exploration–Exploitation Model performs better than the simplified models. The three simplified models tend to capture the mean saccade amplitude rather than the full variety of saccade amplitudes displayed in scan paths. This behavior is expected from the Local Saliency Model, which includes only one type of characteristic saccade amplitude whereas the full Exploration–Exploitation Model has two characteristic saccade amplitudes that correspond either to the exploration or exploitation strategy.

Regarding the Local and Fixed Choice Models, they seem to converge to a behavior where *ρ* is either very small or very big. An extreme value for *ρ* results in a scan path with either exploitation (if *ρ* is close to one) or exploration steps (if *ρ_t_* is close to 0), but almost never both. Hence, these models display a similar behavior to the Local saliency Model which has only one type of characteristic saccade amplitude.

The Bayesian inference process presented in Methods Section results in a distribution over the possible values of the model parameters. This corresponds touncertainty regarding the values of the model parameters. The shading around the generated data curves in Figure5 corresponds to this uncertainty. We sampled 50 different values from the posterior distribution of each one of the model parameters and used this configuration to generate one data set. The shading represents the 95% intervals around the mean density over the different data sets.

The confidence bounds are rather narrow and the density distributions of the two model are highly separable. This is a good indication that the Bayesian parameter inference is reasonable – the saccade amplitude density does not change dramatically with the parameter configurations sampled from the posterior distributions. The confidence bounds for the Local and Fixed Choice Models behave in a similar way and are not included in the figure for clarity purposes.

As described above, we fit a model for each subject individually. Thus, we can investigate how well the Exploration–Exploitation Model captures the difference between the subjects. In Figure6 we compare the mean saccade length of the empirical data and data generated from the Exploration–Exploitation Model for each subject. Each data point is one subject and the diagonal curve is the identity line.

**Fig 6.**
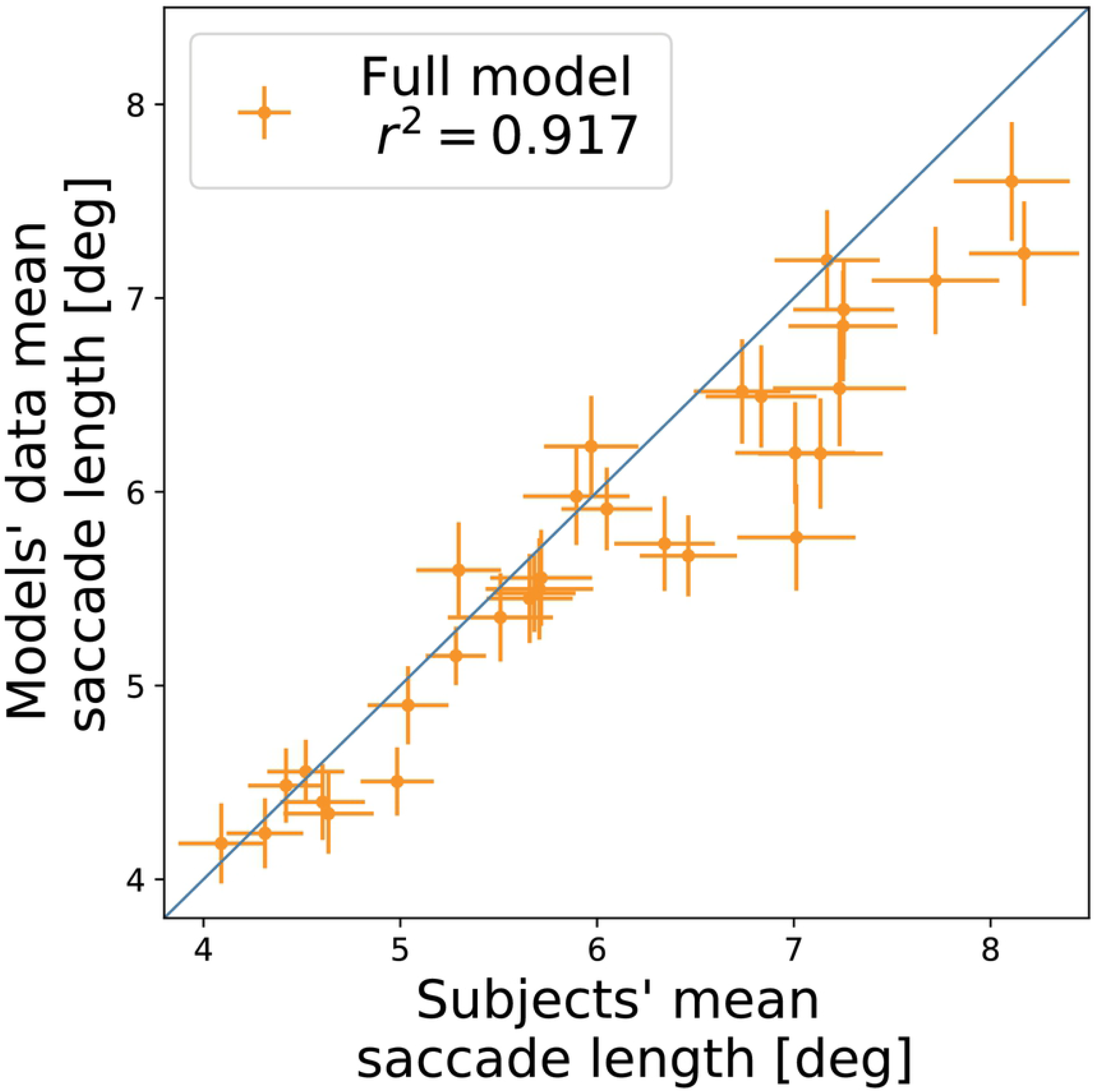
Comparison between experimental and simulated subjects’ mean saccade amplitude. Subjects’ median saccade length compared with the mean saccade length of data generated by the Exploration–Exploitation Model (circles) and by a Local Saliency Model (stars). Overall the Exploration–Exploitation Model captures the median saccade length more accurately than the Local Saliency Model.

Overall, the model captures the different mean saccade length of the different subjects. It seems that the model has a slight tendency to underestimate the mean saccade amplitude, which is also visible in Figure5. The error bars in the figure correspond to the standard error over the different trials for each subject. By comparing the standard error of the empirical saccade amplitudes and of the standard error of the data generated from the Exploration–Exploitation Model, we conclude that the model captures not only the mean saccade amplitude but also the variability of the different saccade amplitudes of each subject.

### Saccade direction

Generally, saccades can be seen as vectors characterized by amplitude and direction. After analyzing the model performance with regard to the saccade amplitude, we turn to analyze the model performance with respect to the saccade direction.

In Figure7 we compare the saccade direction density, over the entire population of subjects, of the empirical data and of data generated by the fitted model. The empirical data demonstrate clear preference for horizontal saccades and a weaker tendency towards vertical saccades. The data generated by the model correspond to a tendency to perform horizontal saccades, but this tendency is not as strong as in the empirical data. The empirical tendency towards vertical saccades is not captured at all by the model.

**Fig 7.**
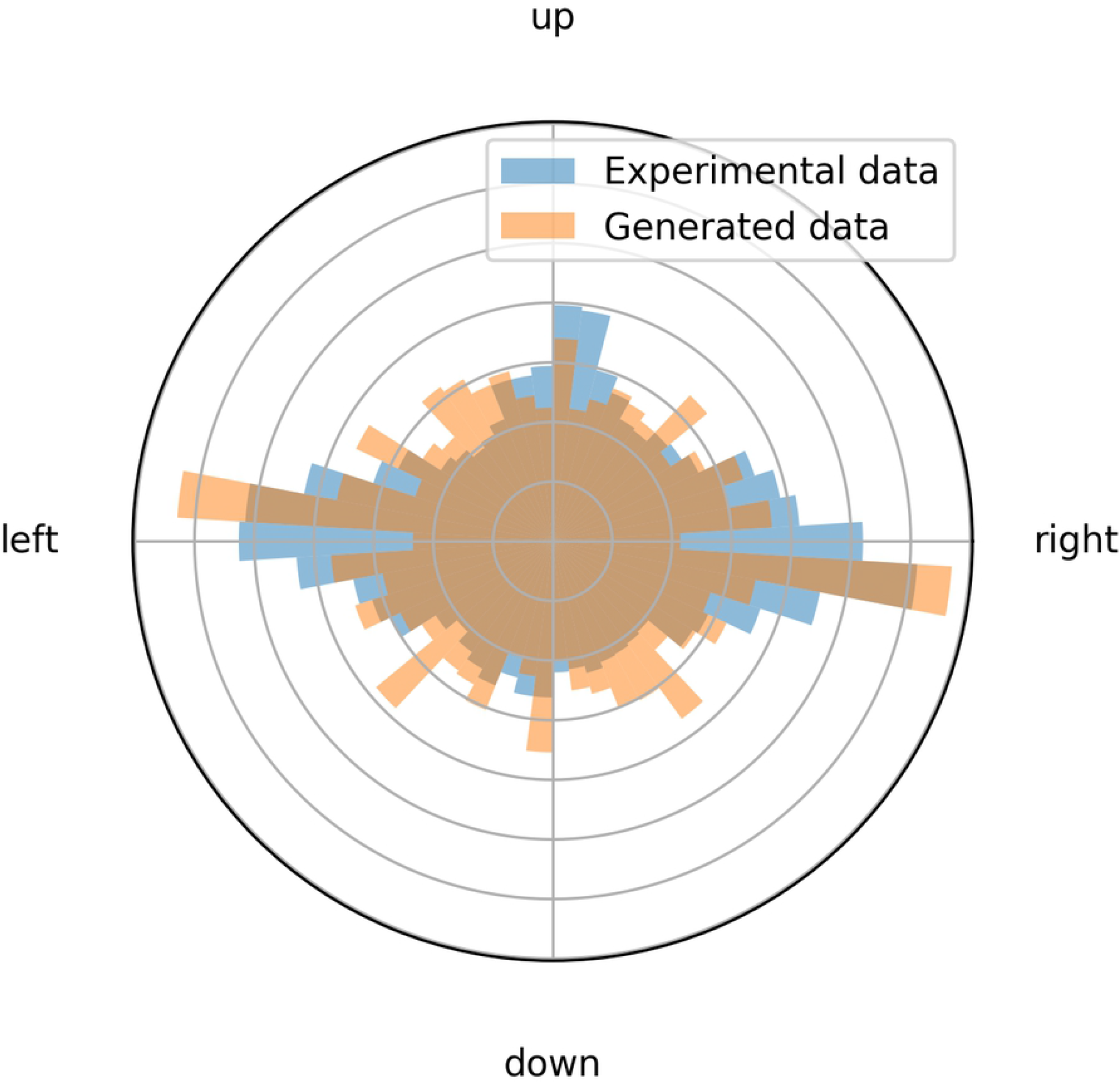
Experimental and simulated saccade direction density. Saccade direction frequency, aggregated over the data from all subjects. In blue is the empirical data and in orange the data generated from the model. The empirical data demonstrate a strong tendency to saccades which are in the left and right directions, and a tendency to perform saccades directed upwards. The model captures the tendency to perform horizontal saccades, but not vertical saccades.

The fact that the model captures only one preferred direction is not surprising. At each step, the next fixation is chosen from a Gaussian distribution in case of an exploitation step, and from a donut–shaped distribution in case of an exploration step. Both of these distributions have only one preferred direction, which results in the oval shape of the saccade direction density. In the Discussion we suggest variations of the model which could capture more than one preferred saccade direction.

### Model comparison

Last, we would like to compare the performance of the full Exploration–Exploitation Model to the simplified variants of the model presented in the Methods Section. As suggested in the work of Kümmerer et al. in [27] we use the information gain to compare the performances of the different models. They define information gain, as the average difference of the log-likelihood between a model and some baseline model. As a baseline we take a uniform distribution over the image, where the log-likelihood for each fixation is constant and equal to log_2_ (number of pixels).

We estimate the likelihood of the different models over different realizations of the model parameters sampled from the posterior. As the model parameters where fitted on the training data, the resulting posterior distribution can be seen as a new prior distribution over the parameters for the test data-set. Thus, the information gain, averaged over different realizations of parameters, is the likelihood ratio test which is a proper scoring rule [28,29].

Table 1 presents of the result of the comparison of the information gain of the different variations of the model with respect to the baseline model. The full Exploration–Exploitation Model has the highest information gain which verifies the assumptions underlying the model over the models without these assumptions. As in Figure5 the uncertainty in the results is over different realizations of the model parameters from the fitted posterior distributions.

**Table 1.** Comparison of the information gain, of the different models measured in bit/fixation. The information gain for each model is computed against a model with a uniform distribution over the image. The Full Exploration–Exploitation Model has the highest information gain which implies it is the most probable model.

|  | Exploration–<br>Exploitation | Local<br>Choice | Fixed<br>Choice | Local<br>Saliency |
| --- | --- | --- | --- | --- |
| Information gain | $2.0 \pm 0.01$ | $1.92 \pm 0.02$ | $1.97 \pm 0.01$ | $1.99 \pm 0.004$ |

## Discussion

The current study proposed and analyzed a mathematical model of fixation selection, motivated by the Exploration–Exploitation paradigm [5]. We constructed a generative scan path model based on a small set of assumption. Using Bayesian [30] inference we fit the model to experimental data. By doing so, we continue the line of work of using generative likelihood based models for scan path generation [18]. Importantly, we use recent developments in Bayesian statistics to construct more efficient parameter inference algorithms.

A different approach uses of deep neural networks for scan path modeling [12, 13]. One of the downsides of this approach is its reliance on large amounts of data, which precludes the study of interindividual differences. Thus, by using a hypothesis-based model, which requires only a relatively small number of parameters, we can fit individual models for each experimental subject and capture inter-subject variability. We demonstrate how our model captures the saccade amplitude both at the population and the individual level.

Another benefit of the Bayesian approach is the ability to easily test different hypotheses. We do so by constructing multiple models, of increasing complexity. Each of the models is fit to the training data and we use a proper scoring rule, namely the likelihood ratio, to compare the different models. This allows us to conclude that the set of hypotheses used for full Exploration–Exploitation Model is more suitable for the describing the experimental data than the reduced models.

As described above, the Exploration–Exploitation Model successfully captures the experimental saccade amplitudes both at the population level and the subject level. Another spatial aspect of saccades is saccade direction. Our model captures only the tendency to perform horizontal saccades, but not the tendency to perform vertical saccades. This is expected from the construction of the model.

In its current form the Exploration–Exploitation Model does not capture the change in saccade direction (i.e. the saccade direction relative to the previous saccade). The relative saccade direction is important for modeling known phenomena such as visual persistence [31, 32]. As with the vertical preferred saccade direction, the model’s inability to capture the relative saccade direction stems from the choice of Gaussian functions in the exploration and exploitation policies. The model can be extend and account for these tendencies by using a mixture model for generating the exploration and exploitation steps. In this case, instead of sampling the next step from a Gaussian distribution in the case of an exploitation step, or a subtraction of Gaussians in the case of exploration step, a mixture of Gaussians could be introduced. Each Gaussian component could be designed to capture different directional tendencies, rather than capturing only one tendency as in the current version of the model.

Other limitations of the model stem from the choice of a second order Markov process. Due to this choice the model is almost memory-less and cannot capture known phenomena in scene viewing which span multiple saccades. One example of such a phenomenon is the well–known inhibition of return [33, 34]. Incorporating longer history is not straightforward in our model. A heuristic approach could be including dynamics in the saliency map. Currently we use a constant saliency map, but the model could be adapted to use an evolving saliency map accounting for effects such as inhibition of return.

Finally, our mathematical model does not account for fixation durations in scene viewing, which play an important role in eye-movement control [17, 35–37]. So far most of the modeling attempts of scene viewing addressed either the spatial or the temporal aspects of scene viewing. Indeed, some models use temporal dynamics but they do not attempt to learn these dynamics from the data and use a heuristic–based approach. While fixation duration modeling is outside the scope of this work, we nonetheless consider the integration of temporal and spatial aspects an exciting new research direction.

## Supporting information

**S1 Appendix. Parameter inference.** In this appendix you can find the technical details of the Gibbs sampler described in the Methods section. It includes a derivation of the full likelihood, details regarding the augmentation schemes and the derivation of the conditional distributions.

## Acknowledgments

This research has been funded by Deutsche Forschungsgemeinschaft (DFG, German Research Foundation) - SFB 1294/1 - 318763901.

## References

1. Chalupa LM, Werner JS. The Visual Neurosciences, Vols. 1 & 2. MIT Press; 2004.

2. Findlay JM, Gilchrist ID. Active Vision: The Psychology of Looking and Seeing. Oxford University Press; 2003.

3. Gameiro RR, Kaspar K, König S, Nordholt S, König P. Exploration and Exploitation in Natural Viewing Behavior. Scientific Reports. 2017;7(1). doi:10.1038/s41598-017-02526-1.

4. Ehinger BV, Kaufhold L, Köonig P. Probing the temporal dynamics of the Exploration–Exploitation dilemma of eye movements. Journal of Vision. 2018;18(3):6. doi:10.1167/18.3.6

5. Berger-Tal O, Nathan J, Meron E, Saltz D. The Exploration–Exploitation dilemma: a multidisciplinary framework. PloS ONE. 2014;9(4):e95693. doi:10.1371/journal.pone.0095693

6. Bisley JW, Mirpour K. The neural instantiation of a priority map. Current Opinion in Psychology. 2019; p. 108–112.

7. Itti L, Koch C. A saliency-based search mechanism for overt and covert shifts of visual attention. Vision Research. 2000;40(10-12):1489–1506.

8. Borji A, Itti L. State-of-the-art in visual attention modeling. IEEE Transactions On Pattern Analysis And Machine Intelligence. 2012;35(1):185–207.

9. Kummerer, Theis, Bethge. Deep Gaze I: Boosting saliency prediction with feature maps trained on imagenet. Preprint arXiv:14111045. 2014.

10. Kummerer, Wallis, Bethge. Deep Gaze II: Reading fixations from deep features trained on object recognition. Preprint arXiv:161001563. 2016.

11. Einhauser W, Konig P. Getting real – sensory processing of natural stimuli. Current Opinion in Neurobiology. 2010;20(3):389–395.

12. Shao X, Luo Y, Zhu D, Li S, Itti L, Lu J. Scanpath prediction based on high-level features and memory bias. In: International Conference on Neural Information Processing. Springer; 2017:3–13.

13. Kummerer M, Wallis TS, Bethge M. DeepGaze III: Using Deep Learning to Probe Interactions Between Scene Content and Scanpath History in Fixation Selection. In: 2019 Conference on Cognitive Computational Neuroscience, 13–.16 September 2019, Berlin, Germany; 2019.

14. Zelinsky GJ. A theory of eye movements during target acquisition. Psychological Review. 2008;115(4):787–835.

15. Le Meur O, Liu Z. Saccadic model of eye movements for free-viewing condition. Vision Research. 2015;116:152–164.

16. Engbert R, Trukenbrod HA, Barthelmóe S, Wichmann FA. Spatial statistics and attentional dynamics in scene viewing. Journal of Vision. 2015;15(1):14. doi:10.1167/15.1.14.

17. Tatler BW, Brockmole JR, Carpenter RH. LATEST: A model of saccadic decisions in space and time. Psychological Review. 2017;124(3):267–300.

18. Schutt HH, Rothkegel LO, Trukenbrod HA, Reich S, Wichmann FA, Engbert R. Likelihood-based Parameter Estimation and Comparison of Dynamical Cognitive Models. Psychological Review. 2017;124(4):505–524.

19. Tatler BW, Baddeley RJ, Vincent BT. The long and the short of it: Spatial statistics at fixation vary with saccade amplitude and task. Vision Research. 2006;46(12):1857–1862.

20. Geman S, Geman D. Stochastic Relaxation, Gibbs Distributions, and the Bayesian Restoration of Images. In: Readings in Computer Vision. Elsevier; 1987:564–584.

21. Liu JS. Monte Carlo strategies in scientific computing. Springer Science & Business Media; 2008.

22. Gelman A, Carlin JB, Stern HS, Dunson DB, Vehtari A, Rubin DB. Bayesian data analysis. Chapman and Hall/CRC; 2013

23. Polson NG, Scott JG, Windle J. Bayesian Inference for Logistic Models Using Pólya-Gamma Latent Variables. Journal of the American Statistical Association. 2013;108(504):1339–1349.

24. Gilks WR, Wild P. Adaptive Rejection Sampling for Gibbs Sampling. Journal of the Royal Statistical Society: Series C (Applied Statistics). 1992;41(2):337–348.

25. Martino L, Yang H, Luengo D, Kanniainen J, Corander J. A fast universal self-tuned sampler within Gibbs sampling. Digital Signal Processing. 2015;47:68–83.

26. Duane S, Kennedy AD, Pendleton BJ, Roweth D. Hybrid Monte Carlo. Physics Letters B. 1987;195(2):216–222.

27. Kümmerer M, Wallis TS, Bethge M. Information-theoretic model comparison unifies saliency metrics. Proceedings of the National Academy of Sciences. 2015;112(52):16054–16059.

28. Cox DR. Tests of separate families of hypotheses. Proceedings of the fourth Berkeley symposium on mathematical statistics and probability. 1961;1:105–123.

29. Lewis F, Butler A, Gilbert L. A unified approach to model selection using the likelihood ratio test. Methods in Ecology and Evolution. 2011;2(2):155–162.

30. MacKay DJ, Mac Kay DJ. Information theory, inference and learning algorithms. Cambridge University Press; 2003.

31. Ritter M. Evidence for visual persistence during saccadic eye movements. Psychological Research. 1976;39(1):67–85.

32. Breitmeyer BG, Kropfl W, Julesz B. The existence and role of retinotopic and spatiotopic forms of visual persistence. Acta Psychologica. 1982;52(3):175–196.

33. Handy TC, Jha AP, Mangun GR. Promoting novelty in vision: Inhibition of return modulates perceptual-level processing. Psychological Science. 1999;10(2):157–161.

34. Klein RM. Inhibition of return. Trends in Cognitive Sciences. 2000;4(4):138–147.

35. Henderson JM. Human gaze control during real-world scene perception. Trends in Cognitive Sciences. 2003;7(11):498–504.

36. Nuthmann A, Smith TJ, Engbert R, Henderson JM. CRISP: a computational model of fixation durations in scene viewing. Psychological Review. 2010;117(2):382–405.

37. Laubrock J, Cajar A, Engbert R. Control of fixation duration during scene viewing by interaction of foveal and peripheral processing. Journal of Vision. 2013;13(12). doi:10.1167/13.12.11.

